# Cdc42 GTPase activating proteins Rga4 and Rga6 coordinate septum synthesis and membrane trafficking at the division plane during cytokinesis

**DOI:** 10.1101/2021.12.14.472679

**Authors:** Bethany F. Campbell, Brian S. Hercyk, Ashlei R. Williams, Ema San Miguel, Haylee G. Young, Maitreyi E. Das

## Abstract

Fission yeast cytokinesis is driven by simultaneous septum synthesis, membrane furrowing and actomyosin ring constriction. The septum consists of a primary septum flanked by secondary septa. First, delivery of the glucan synthase Bgs1 and membrane vesicles initiate primary septum synthesis and furrowing. Next, Bgs4 is delivered for secondary septum formation. It is unclear how septum synthesis is coordinated with membrane furrowing. Cdc42 promotes delivery of Bgs1 but not Bgs4. We find that after primary septum initiation, Cdc42 inactivators Rga4 and Rga6 localize to the division site. In *rga4Δrga6Δ* mutants Cdc42 activity is enhanced during late cytokinesis and cells take longer to separate. Electron micrographs of the division site in these mutants exhibit malformed septum with irregular membrane structures. These mutants have a larger division plane with enhanced Bgs1 delivery but fail to enhance accumulation of Bgs4 and several exocytic proteins. Additionally, these mutants show endocytic defects at the division site. This suggests that Cdc42 regulates only specific membrane trafficking events. Our data indicate that while active Cdc42 promotes primary septum synthesis, as cytokinesis progresses Rga4 and Rga6 localize to the division site to decrease Cdc42 activity. This couples specific membrane trafficking events with septum formation to allow proper septum morphology.

**Abstract Figure:** 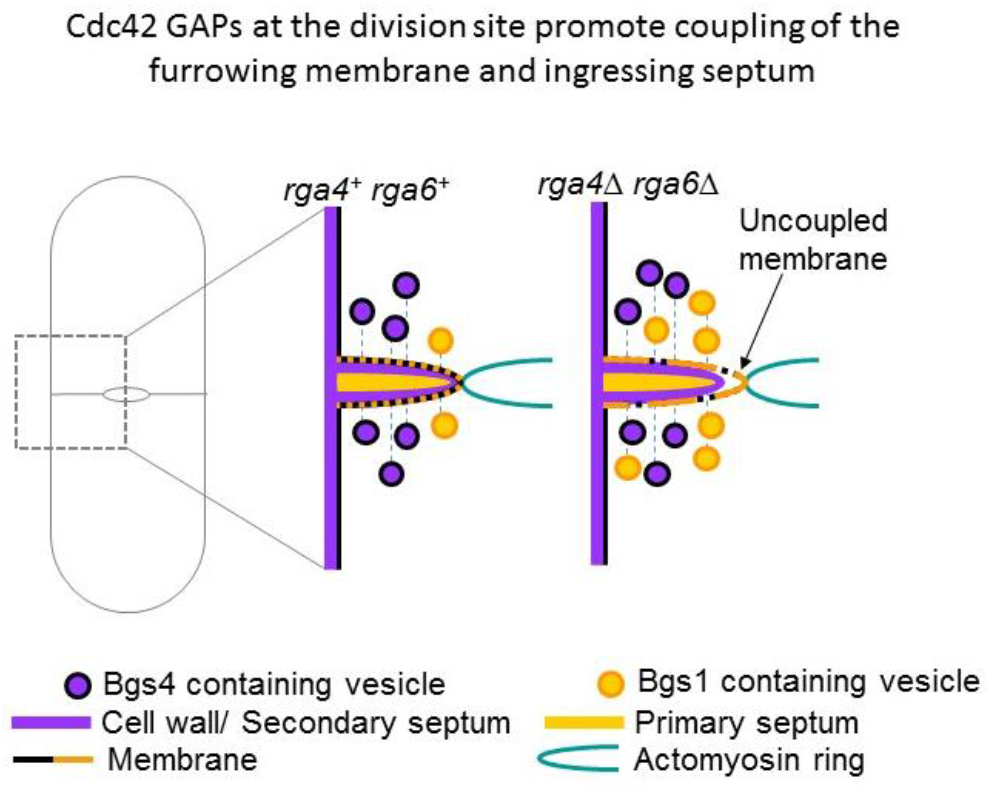

**Synopsis:** The GTPase Cdc42 regulates cytokinesis in cell-walled fission yeast. Active Cdc42 promotes the initiation of septum (new cell wall) synthesis to physically divide daughter cells. Here we show that Cdc42 activity must be decreased at the later stages of cytokinesis to enable proper septum formation. Mutants lacking Cdc42 inactivators, Rga4 and Rga6, lead to membrane trafficking defects and malformed septa consequently delaying cell separation.

## Introduction

Cytokinesis, the final step in cell division is a complex process involving multiple events. In fission yeast *Schizosaccharomyces pombe*, cytokinesis involves the assembly of an actomyosin ring which constricts along with membrane furrow formation and septum (cell wall) synthesis. It is unclear how ring constriction is tightly coupled to membrane furrowing and septum formation. Cdc42 is a central regulator of cell polarity and cell growth in most eukaryotes ^1^. Recent work, however, has demonstrated that Cdc42 is also required for certain steps of cytokinesis in *S. pombe* ^2,3^. During cytokinesis, numerous proteins are delivered in a sequential manner via distinct membrane trafficking events to ensure the fidelity of division. First, proteins necessary to form the contractile actomyosin ring are recruited to cortical nodes in the cell middle where division will occur ^4^. After the actomyosin ring assembles and matures, septum-synthesizing enzymes are delivered to build the tri-layer division septum that consists of a primary septum flanked by two secondary septa ^5^. In addition, membrane is also delivered to the division site for furrow formation ^6^. Once the septum matures, the primary septum is digested to separate the daughter cells, while the secondary septa remain as daughter cell wall. In *S. pombe*, the primary septum is mostly synthesized by the linear-β-glucan synthase Bgs1 ^7^, while the secondary septa are primarily synthesized by the branched β-glucan synthase Bgs4 ^7,8^. Once Bgs1 arrives at the membrane adjacent to the actomyosin ring, primary septum synthesis initiates to drive ring constriction ^2,5,9,10^. Shortly after septum synthesis initiation, Bgs4 is delivered to the division site to synthesize the secondary septa which further promotes and stabilizes ring constriction ^8^. Finally, after completion of ring constriction and septum maturation, the primary layer of the septum is digested by the glucanases Eng1 and Agn1 to physically separate the daughter cells, completing division ^11-16^. Cdc42 activity is necessary for the delivery of the primary septum synthase Bgs1 as well as the septum digesting glucanases Eng1 and Agn1 ^2,3^.

In addition to temporally ordering the events of cytokinesis to ensure proper ring formation, ring constriction, membrane furrowing, and septum synthesis, membrane trafficking events are also spatially regulated to ensure that cargo is properly delivered to and removed from specific sites along the division plane^17^. In *S. pombe*, endocytosis occurs primarily at the outer rim of the membrane furrow, while exocytosis occurs all along the division plane to deliver protein cargo and new membrane to the division site ^6^. Specifically, the exocyst complex tethers exocytic vesicles at the outer edge of the membrane furrow, while the TRAPP-II complex regulates exocytosis throughout the whole membrane furrow to promote the tethering of vesicles containing the Rab11 GTPase Ypt3 and to a lesser extent the Rab8 GTPase Ypt2 ^6^. Notably, the TRAPP-II complex promotes the delivery of both the primary septum synthase Bgs1 as well as the secondary septum synthase Bgs4 to the division site ^6^. To ensure efficient delivery of exocytic vesicles carrying cargo necessary for cytokinesis, a type V myosin motor Myo52 physically transports many of these vesicles along actin cables to the division site, although they can also arrive via random walk ^6,18^. Thus, cytokinetic membrane trafficking events are spatiotemporally regulated to control the delivery and removal of cargo and thereby coordinate ring constriction, membrane ingression, and septum synthesis.

Notably, Cdc42 activity is itself spatiotemporally regulated at the division plane via Guanine nucleotide Exchange Factors (GEFs) Gef1 and Scd1 which differentially regulate its activity ^2,19^. During cytokinesis, Gef1 localizes to the cytokinetic ring and activates Cdc42 immediately after ring assembly, while Scd1 localizes behind the already constricting ring and activates Cdc42 along the ingressing membrane furrow ^2^. Cdc42 is inactivated by the GTPase activating proteins (GAPs) Rga4, Rga6, and Rga3 ^20-23^. These GAPs have been shown to localize to the division site, but their role in cytokinesis is unknown ^20-23^. Rga4 and Rga6 are the primary and secondary Cdc42 GAPs, respectively, since *rga4Δ* cells are wider than *rga6Δ* cells, while *rga3Δ* cells exhibit no morphology changes ^20,22,23^. The primary Cdc42 GAP Rga4 localizes to the division site after the initiation of septum synthesis ^20^. Similarly, the secondary Cdc42 GAP Rga6 localizes to the division site long after Cdc42 is activated at the division plane ^22^. The third Cdc42 GAP Rga3 appears to function redundantly with the other two GAPs during vegetative growth, however, it is also present at the division site where it localizes in a manner similar to Cdc42 ^23^. Thus, it seems that Cdc42 is activated at the division site to initiate primary septum synthesis and drive cytokinetic ring constriction, however, it is unclear what role the GAPs play at the division site during cytokinesis.

Here we show that the GAPs Rga4 and Rga6 decrease Cdc42 activity after septum formation initiates to coordinate septum synthesis and membrane trafficking within the division plane. In the absence of these GAPs, the balance between different membrane trafficking events is perturbed, spatiotemporal pattern of endocytosis is disrupted, septa and membrane furrow exhibit morphological defects, and cell separation is delayed. These findings indicate that the spatiotemporal Cdc42 activation pattern is required to couple membrane furrowing with septum synthesis during cytokinesis.

## Results

### The GAPs Rga4 and Rga6 localize to the membrane furrow to decrease Cdc42 activity during actomyosin ring constriction

Cdc42 is activated at the division site during actomyosin ring assembly ^2,19,24^. While each Cdc42 GAP also localizes to the division site, their localization patterns throughout cytokinesis have not been examined in detail. We observed the localizations of the GAPs relative to the actomyosin ring throughout each stage of cytokinesis. We find that Rga4-GFP and Rga6-GFP do not localize to the assembled actomyosin ring as marked by Rlc1-tdTomato before the onset of constriction (Figure 1A and B). In contrast, Rga3-GFP localizes to the assembled ring before constriction initiates (Figure 1C). Rga4-GFP and Rga6-GFP instead localize to the ingressing membrane after the ring initiates constriction. Additionally, Rga4-GFP and Rga6-GFP remain at the division site following completion of ring constriction and loss of Rlc1-tdTom signal (Figure 1A and B). Rga3-GFP signal is lost from the division site following ring constriction unlike that of Rga4-GFP and Rga6-GFP (Figure 1C). Previous reports do not show an important role for Rga3 in regulating Cdc42 during vegetative growth ^23^. In addition, we find that Rga3-GFP localizes to the division site during early cytokinesis, when Cdc42 activity at this site is high and is lost from the division site once ring constriction ends. In contrast, Rga4 and Rga6 localize to the division site only after ring constriction and septum synthesis initiates and remain at this site throughout cytokinesis. Therefore, we investigated the role of Rga4 and Rga6 in late cytokinesis.

**Figure 1.**
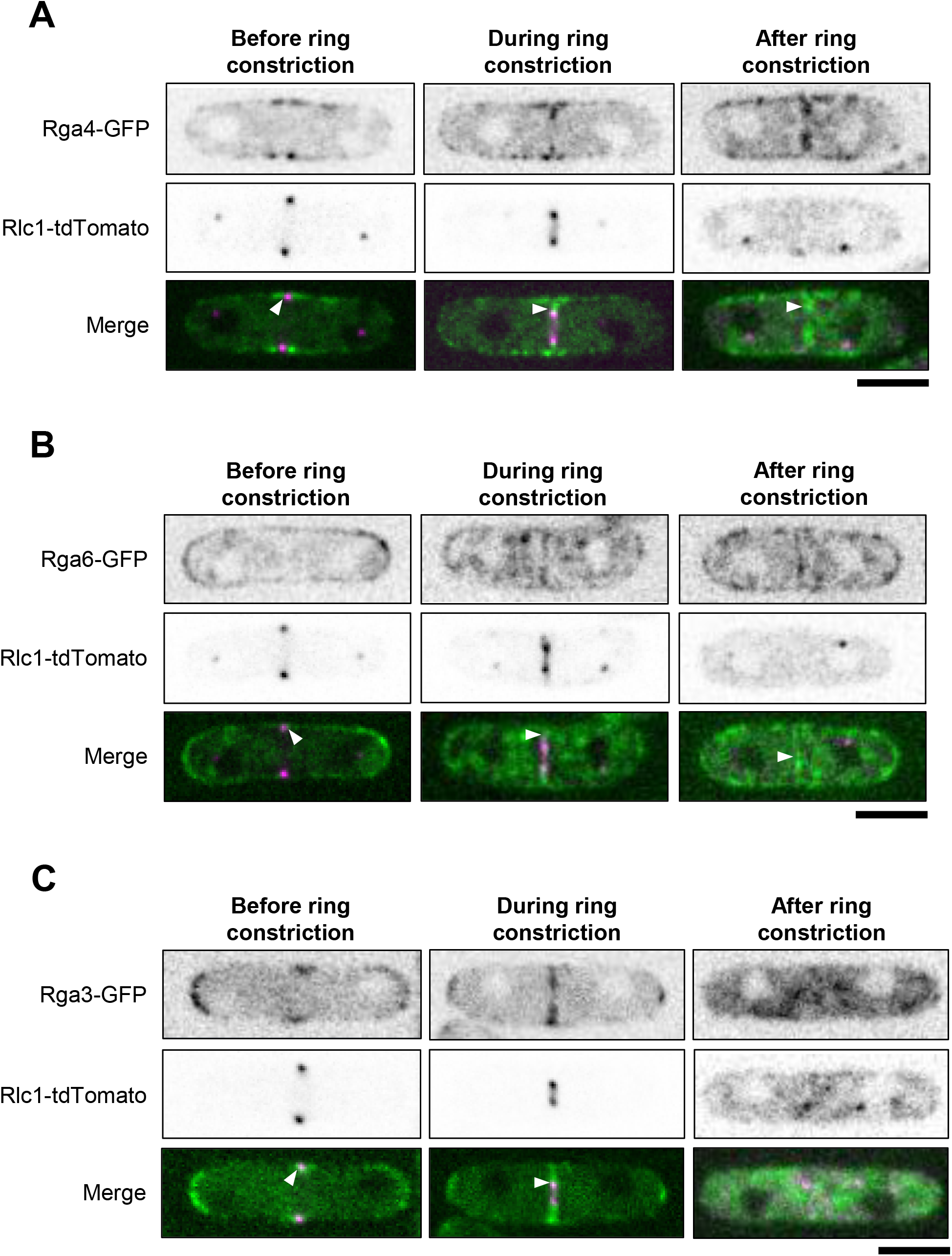
The GAPs Rga4 and Rga6 localize to the ingressing membrane furrow during cytokinesis. Before the onset of ring constriction, Rga4-GFP (A) and Rga6-GFP (B) do not co-localize with the ring marker Rlc1-tdTom (white arrowheads). During ring constriction, Rga4-GFP (A) and Rga6-GFP (B) localize to the membrane furrow just behind the actomyosin ring where they spread through the membrane furrow. After completion of ring constriction, both Rga4-GFP (A) and Rga6-GFP (B) remain at the division site (white arrowheads). (C) Rga3 co-localizes with the actomyosin ring both before and during ring constriction (white arrowheads). After ring constriction, Rga3-GFP is lost from the division site (white arrowhead). scale bar = 5μm

Since Cdc42 is first activated at the division site during ring assembly ^2,19,24^ while the GAPs Rga4 and Rga6 localize to the division site during ring constriction, we reasoned that these GAPs locally decrease Cdc42 activity at the division site during late cytokinesis. To test this, we deleted the GAP genes *rga4* and *rga6* and measured Cdc42 activation levels during late ring constriction in *rga4Δrga6Δ* cells compared to *rga4*^*+*^*rga6*^*+*^. Using a bio-probe that specifically binds active Cdc42 (CRIB-3xGFP) ^21^, we observed that Cdc42 activation at the division site is increased in the absence of *rga4* and *rga6* (Figure 2 A-C). During late ring constriction, the sum CRIB-3xGFP intensity is increased in cells lacking *rga4* and *rga6* (Figure 2 A and B). However, *rga4Δrga6Δ* mutant cells are wider than *rga4*^*+*^*rga6*^*+*^ cells (Figure S1A). Thus, it is possible that the increased sum intensity is simply attributable to the wider cell division site. To rule out this possibility, we measured the mean intensity of CRIB-3xGFP at the division site of *rga4Δrga6Δ* mutants and *rga4*^*+*^*rga6*^*+*^ cells. We find that the mean intensity of CRIB-3xGFP at the division site is also higher in *rga4Δrga6Δ* mutant cells as compared to *rga4*^*+*^*rga6*^*+*^ cells (Figure 2C). These findings indicate that both the total amount of active Cdc42 and the average Cdc42 signal are increased at the division site in the absence of these GAPs.

**Figure 2.**
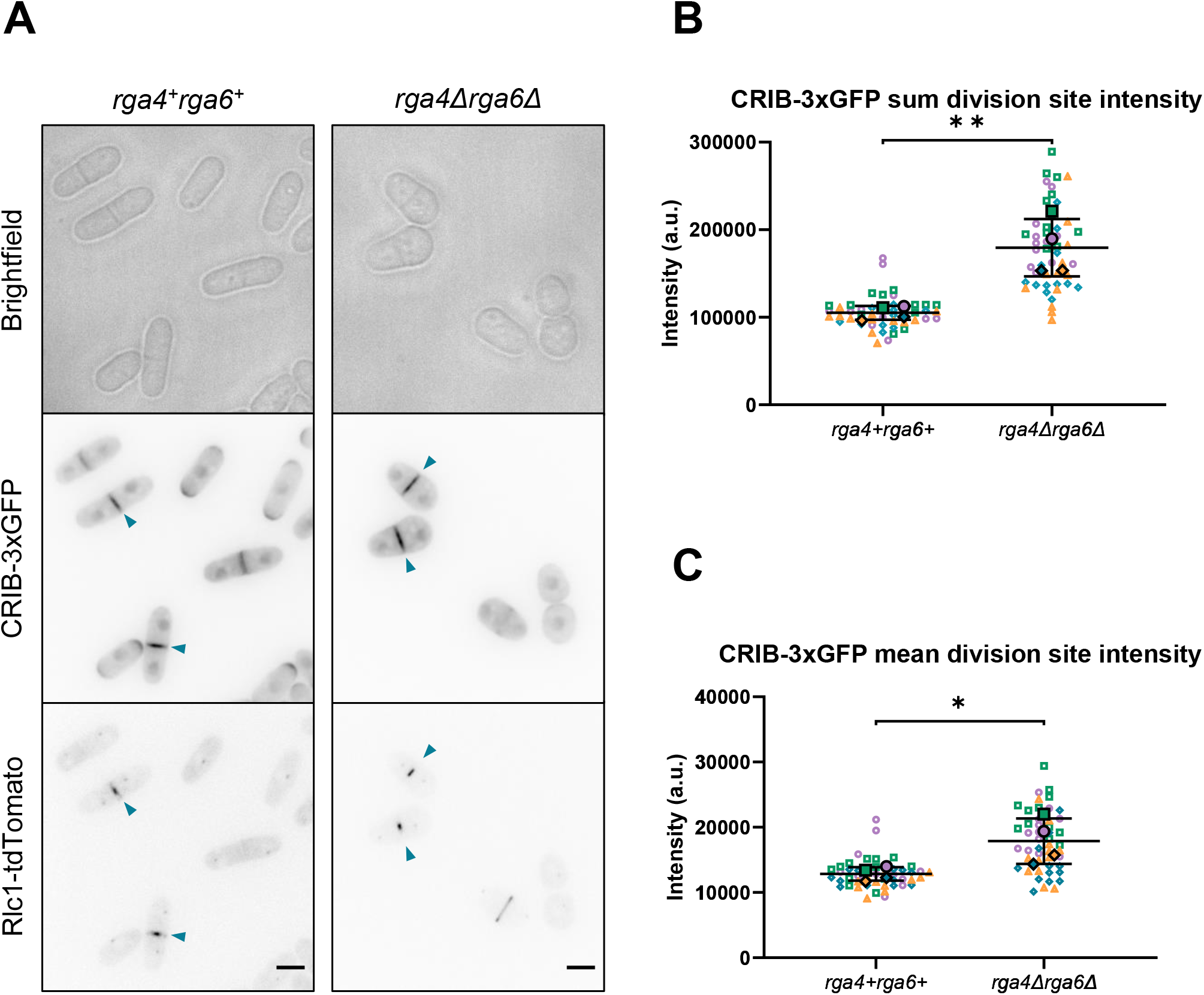
Cdc42 activation at the division site is increased during ring constriction in the absence of *rga4* and *rga6*. (A) Cdc42 activity marked with the bio-probe CRIB-3xGFP is enhanced at the division site (blue arrow heads) in *rga4Δrga6Δ* cells compared to *rga4*^*+*^*rga6*^*+*^ cells (scale bar = 5 μm). (B) Total CRIB-3xGFP intensity at the division site during ring constriction is increased in the absence of *rga4* and *rga6*. (C) Mean CRIB-3xGFP intensity at the division site during ring constriction is increased in the absence of *rga4* and *rga6*. Colors = distinct experiment. Open symbols = individual cells. Solid symbols = means of each experiment. n≥12 cells per genotype per experiment. Unpaired Student’s *t*-test was used to calculate the statistical significance using the means of each experiment. *, *p*<0.05; **, *p*<0.01.

### The GAPs Rga4 and Rga6 ensure timely cell separation following actomyosin ring constriction

Mutant cells lacking *rga4* and *rga6* show altered cell polarity, resulting in shorter, wider cells (Figure S1A ^22,23^). We wondered if in addition to cell polarization, these GAPs also play a role in cytokinesis. We find that in *rga4Δrga6Δ* cells, the onset of ring constriction is not delayed with respect to *rga4*^*+*^*rga6*^*+*^, *rga4Δ*, and *rga6Δ* cells (Figure 3A). Next, we analyzed ring constriction in these cells. We find that the duration of ring constriction is longer in *rga4Δrga6Δ* mutants compared to *rga4*^*+*^*rga6*^*+*^ (Figure S1B). Since *rga4Δrga6Δ* cells are wider than *rga4*^*+*^*rga6*^*+*^ cells, it is not surprising that the ring takes longer to constrict as this has been observed in other wide mutants such as *scd1Δ* ^2^. However, the rate of ring constriction remains the same in the double GAP mutant compared to the controls, suggesting that there is no defect in constriction (Figure 3B). In contrast, we find that *rga4Δrga6Δ* cells take nearly twice as long to undergo cell separation following completion of ring constriction compared to *rga4*^*+*^*rga6*^*+*^ cells (Figure 3C-E). The single GAP mutants also exhibit a delay in cell separation, but the delay is longer in the double mutant (Figure 3E). These findings suggest that Rga4 and Rga6 function additively to regulate timing of cell separation following completion of ring constriction.

**Figure 3.**
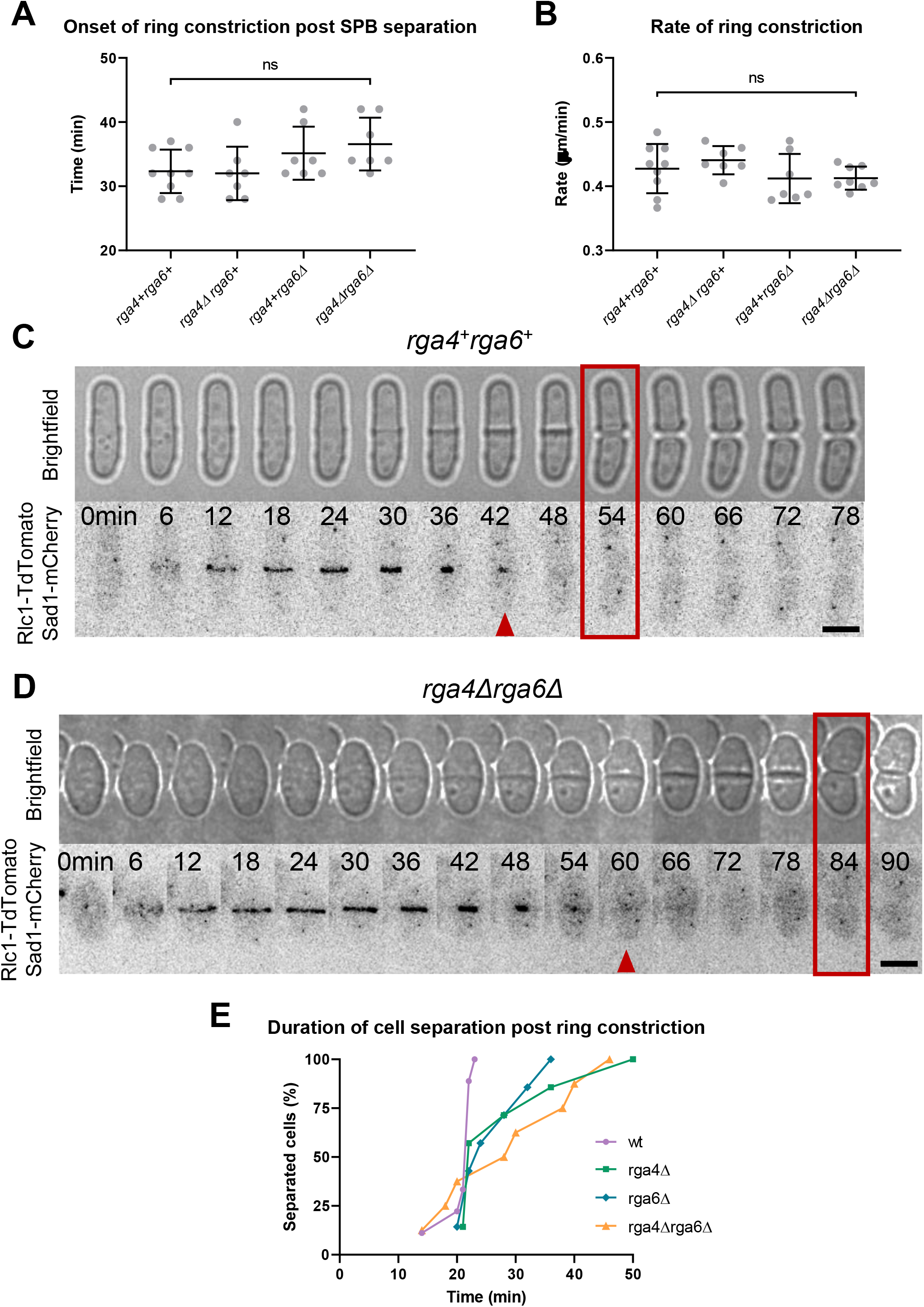
Loss of *rga4* and *rga6* results in delayed cell separation. (A) Onset of ring constriction measured from spindle pole body (SPB) separation is not delayed in the absence of *rga4* and *rga6* (n≥7 cells). (B) Rate of ring constriction remains constant in the absence of *rga4* and *rga6* (n≥7 cells). (C) Example *rga4*^*+*^*rga6*^*+*^ cell separates into two daughter cells (red box) 12 minutes after completion of ring constriction (red arrowhead) where Rlc1-tdTom marks the actomyosin ring. (D) Example *rga4Δrga6Δ* cell takes 24 minutes to separate into daughter cells (red box) following completion of ring constriction (red arrowhead). (E) Individual loss of each GAP in *rga4Δ* and *rga6Δ* cells increases duration of cell separation following completion of ring constriction. The delay in cell separation is more pronounced in the *rga4Δrga6Δ* mutant, where 50% of cells have not separated at 30 minutes post ring constriction, while all *rga4*^*+*^*rga6*^*+*^ cells have successfully separated (n≥7 cells). Statistical test used is Ordinary one-way ANOVA with Tukey’s multiple comparisons. n.s., not significant; scale bar = 5μm.

### Rga4 and Rga6 are required for proper septum morphology

In fission yeast, cell separation delays are often caused by improper localization of the primary septum digesting glucanases Eng1 and Agn1 ^13,14,16^. To probe the cause of delayed cell separation, we examined the localization patterns of Eng1-GFP and Agn1-GFP ^11,12^ at the division site of *rga4Δrga6Δ* mutants compared to *rga4*^*+*^*rga6*^*+*^ cells. Eng1-GFP and Agn1-GFP appear to properly localize to the outer rim of the membrane furrow and to the center of the division plane as a dot in a manner similar to *rga4*^*+*^*rga6*^*+*^ cells (Figure S2A-C ^16^). Since Eng1 and Agn1 are properly delivered to the division site in the absence of *rga4* and *rga6*, the delay in cell separation does not seem to result from the inability of the glucanases to digest the primary septum to physically separate the daughter cells.

Cell separation delays can also occur if the primary septum does not form properly ^7^. Thus, we assessed the structural integrity of the septum in the *rga4Δrga6Δ* mutant cells. To test this, we used transmission electron microscopy (TEM) to image the division sites of *rga4*^*+*^*rga6*^*+*^ and *rga4Δrga6Δ* cells. Wild-type *rga4*^*+*^*rga6*^*+*^ cells typically build straight septa tightly coupled to the membrane furrow (Figure 4A, *i* and *ii*). In contrast, in *rga4Δrga6Δ* cells, several morphological defects were observed in the septa and the plasma membrane flanking the septa (Figure 4A and B). The septa of *rga4Δrga6Δ* cells are spatially disorganized. The leading edge of the ingressing septum did not clearly show the plasma membrane in the mutant cells (Figure 4B, *i* and *ii*, arrowheads). Additionally, some *rga4Δrga6Δ* septa appeared to have plasma membrane trapped within it (Figure 4B, iii). We also found clusters of spherical vesicle-like structures near the synthesizing septa in *rga4Δrga6Δ* cells (Figure 4B, *i, ii*, and *iv*). These structures have a diameter of ∼40nm, but their identity remains to be determined. Thus overall, the division plane in *rga4Δrga6Δ* mutants display morphological defects particularly with respect to membrane organization.

**Figure 4.**
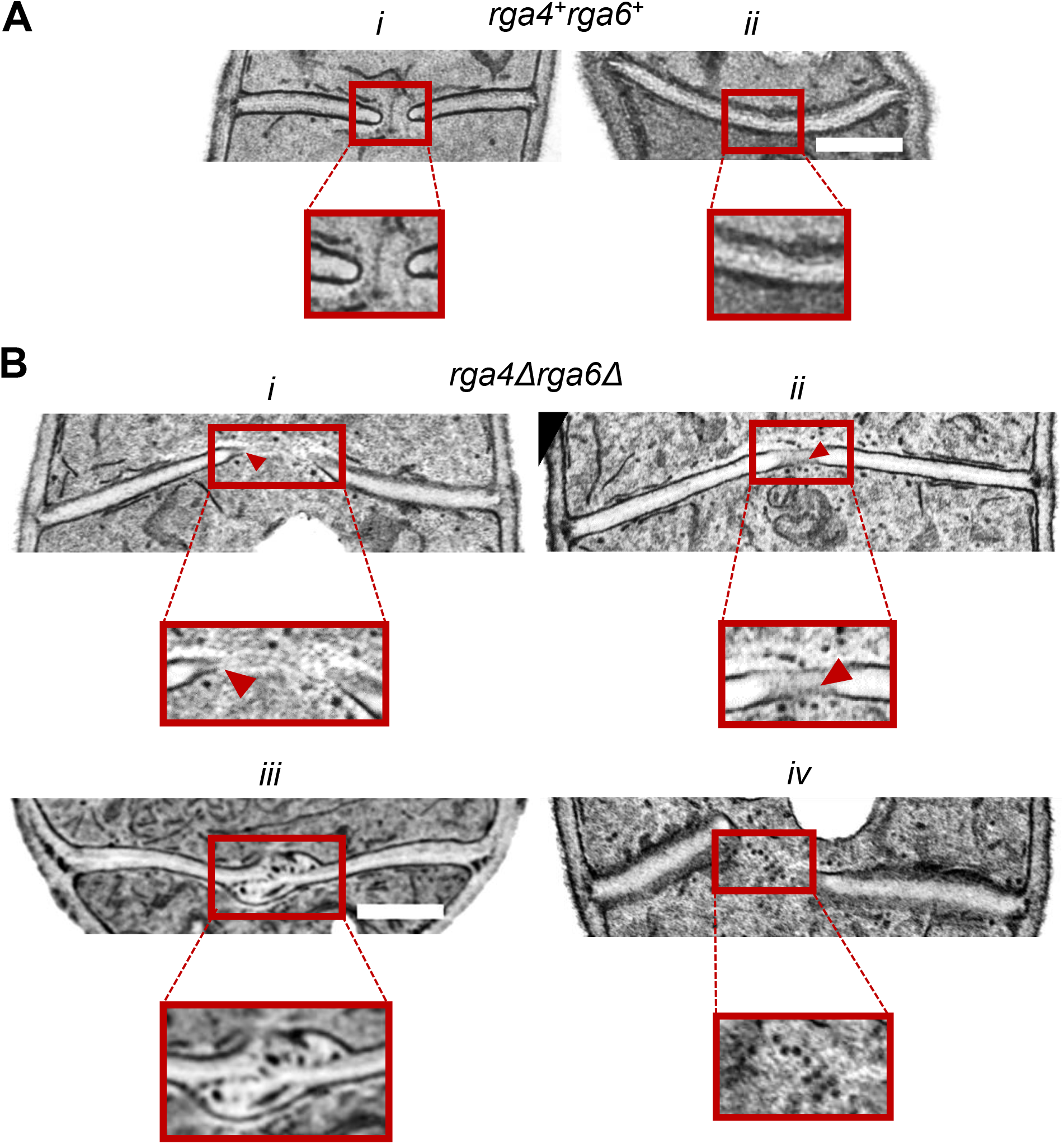
Loss of *rga4* and *rga6* results in malformed division septa. (A) In *rga4*^*+*^*rga6*^*+*^ cells, the plasma membrane (black) flanking the septum (gray) is tightly coupled with the synthesizing septum (*i*) as well as the completed septum (*ii*). (B) In *rga4Δrga6Δ* cells, several septum morphology defects are evident, ranging from plasma membrane that is discontinuously visible along the septum (*i* and *ii*, red arrowheads) to plasma membrane trapped within the completed septum (*iii*). Additionally, an increase in vesicle-like structures near the synthesizing septum was observed (*i, ii, iv*). Scale bar = 1μm

### Rga4 and Rga6 help to maintain the balance of different septum synthesizing enzymes recruited to the division site

Next, we asked if the septum morphology defects observed by TEM were due to the uncoupling or imbalance of septum synthesis and membrane furrowing in these mutants. To test this, we investigated the localization of the septum synthases Bgs1 and Bgs4, which are mainly required for the primary and secondary septa, respectively. Previous reports have shown that Cdc42 is required for the recruitment of Bgs1 to the division site but not that of Bgs4 ^2,3,25^. We measured the intensities of GFP-Bgs1 and GFP-Bgs4 at the division sites of cells during late ring constriction, since this is the stage at which both the primary and the secondary septum are simultaneously synthesized behind the constricting ring. Overall, both synthases localize properly to the division site during late ring constriction in the absence of *rga4* and *rga6* (Figure 5A and D, arrowheads). 3D-reconstructions of division sites in *rga4*^*+*^*rga6*^*+*^ and *rga4Δrga6Δ* cells show that GFP-Bgs1 (Figure 5B) and GFP-Bgs4 (Figure 5E) properly flank constricting and constricted rings marked with Rlc1-tdTom.

**Figure 5.**
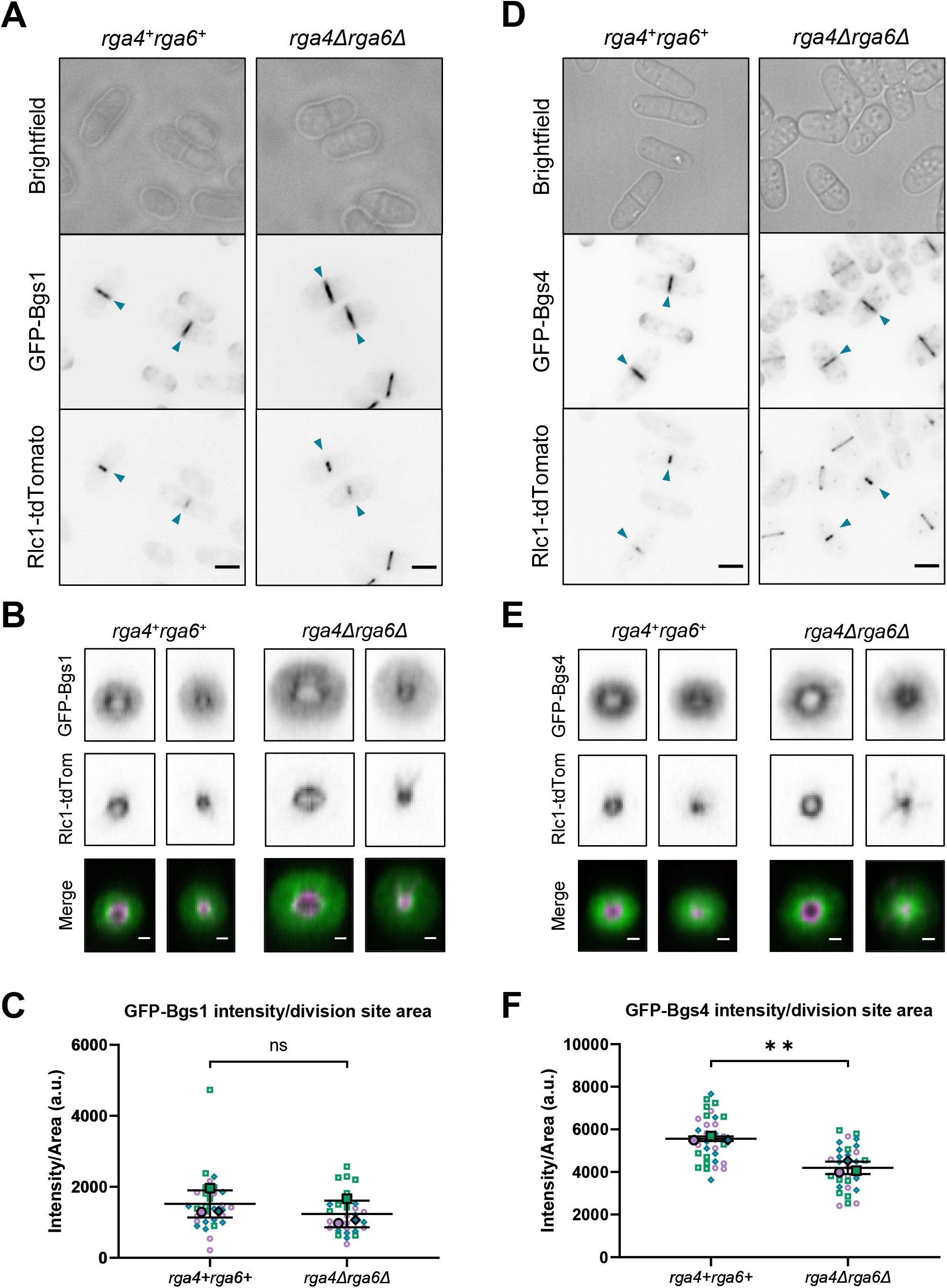
Cdc42-dependent Bgs1 delivery to the division site is enhanced in the absence of *rga4* and *rga6*, while Cdc42-independent Bgs4 delivery is not. (A) GFP-Bgs1 intensity at the division site in *rga4Δrga6Δ* cells is increased compared to *rga4*^*+*^*rga6*^*+*^ cells (scale bar = 5 μm). (B) 3D-reconstruction of the division plane (scale bars = 1 μm) during ring constriction (Rlc1-tdTomato) do not show any defect in the spatial organization of GFP-Bgs1 at the membrane furrow in *rga4Δrga6Δ* cells. (C) GFP-Bgs1 intensity normalized to division site area is not different between *rga4*^*+*^*rga6*^*+*^ and *rga4Δrga6Δ* cells, suggesting that in these mutants Bgs1 delivery enhances along with the enlarged division site. (D) GFP-Bgs4 intensity at the division site in *rga4Δrga6Δ* cells is diminished compared to *rga4*^*+*^*rga6*^*+*^ cells (scale bar = 5 μm). (E) 3D-reconstruction of the division plane during ring constriction (Rlc1-tdTomato) shows decreased intensity of GFP-Bgs4 at the membrane furrow in *rga4Δrga6Δ* cells. (F) GFP-Bgs4 intensity normalized to division site area is less in *rga4Δrga6Δ* cells compared to *rga4*^*+*^*rga6*^*+*^ cells indicating that Bgs4 delivery does not scale with the enlarged division site in these mutants. Colors = distinct experiment. Open symbols = individual cells. Solid symbols = means of each experiment. n≥10 cells per genotype per experiment. Unpaired Student’s *t*-test, was used to calculate statistical significance using the means of each experiment. n.s., not significant; **, *p*<0.01.

As shown previously, *rga4Δrga6Δ* cells are wider than *rga4*^*+*^*rga6*^*+*^ cells and therefore have a larger division site. This suggests that these mutants should have an increased need for the septum synthesizing enzymes due to the enlarged division site. To test this, we measured the intensity of GFP-Bgs1 and GFP-Bgs4 normalized to the area of the division site. We calculated the area of the division site in individual cells and divided the measured sum intensity for each cell by its calculated area. After normalizing the measured intensity for division site area in individual cells, we found that GFP-Bgs1 signal scales with increasing cell width since the overall GFP-Bgs1 sum intensity over area is not altered in the absence of *rga4* and *rga6* (Figure 5C). In contrast, GFP-Bgs4 signal does not scale with increasing cell width since *rga4Δrga6Δ* cells have less GFP-Bgs4 intensity over area than *rga4*^*+*^*rga6*^*+*^ cells (Figure 5F). These findings indicate that loss of *rga4* and *rga6* differentially impacts delivery of the septum synthases Bgs1 and Bgs4. While Cdc42-dependent delivery of Bgs1 scales with increased division site area in the wider *rga4Δrga6Δ* mutant, the amount of Cdc42-independent delivery of Bgs4 to the division site in these cells does not scale. Thus, loss of *rga4* and *rga6* and the resulting increase in Cdc42 activity lead to an enhancement of Cdc42-mediated delivery of Bgs1, while Cdc42-independent delivery of Bgs4 is unaltered.

### Accumulation of exocytic proteins does not scale with enlarged division site in the absence of *rga4* and *rga6*

The septum synthases are known to be delivered to the division site by membrane trafficking events ^6,26-28^. Given the differential recruitment of the septum synthases at the division site in *rga4Δrga6Δ* mutants we asked if membrane trafficking was correspondingly altered in these mutants. Previous reports suggest that both Bgs1 and Bgs4 are delivered via vesicle trafficking pathways that include the Ypt3 (Rab11 GTPase), Trs120 (a component of the TRAPP-II complex), Syb1 (a v-SNARE), and Myo52 (a Type V myosin motor) ^6,27^. Thus, we measured the levels of these vesicular trafficking markers at the division site. Following the same paradigm used with the septum synthases, we measured the sum intensities normalized to the division site area for mEGFP-Ypt3, Trs120-3xGFP, GFP-Syb1, and Myo52-GFP. We specifically chose to examine the localization of Ypt3 since it has been shown to be involved in TRAPP-II-mediated exocytosis throughout the membrane furrow and since its puncta largely co-localize with the septum synthase Bgs4, a vesicle cargo ^6^. When mEGFP-Ypt3 signal is normalized to the area of the division site, there appears to be less Ypt3 accumulation in the absence of *rga4* and *rga6* (Figure 6B). The overall distribution of Ypt3 within the division plane, however, is unperturbed in *rga4Δrga6Δ* cells compared to *rga4*^*+*^*rga6*^*+*^ cells indicating that Ypt3 likely remains functional, albeit is less abundant relative to the size of the division site (Figure 6A). Next, we examined the localization of Trs120, a subunit of the TRAPP-II complex which largely co-localizes with Ypt3 to promote exocytic vesicle tethering and which promotes delivery of Bgs4 puncta, both at the division site and the cell ends ^6^. After normalizing to the size of the division site, Trs120-3xGFP levels do not scale to division site area in the absence of *rga4* and *rga6* (Figure 6D). We did not see any change in the localization pattern of Trs120 at the division site (Figure 6C).

**Figure 6.**
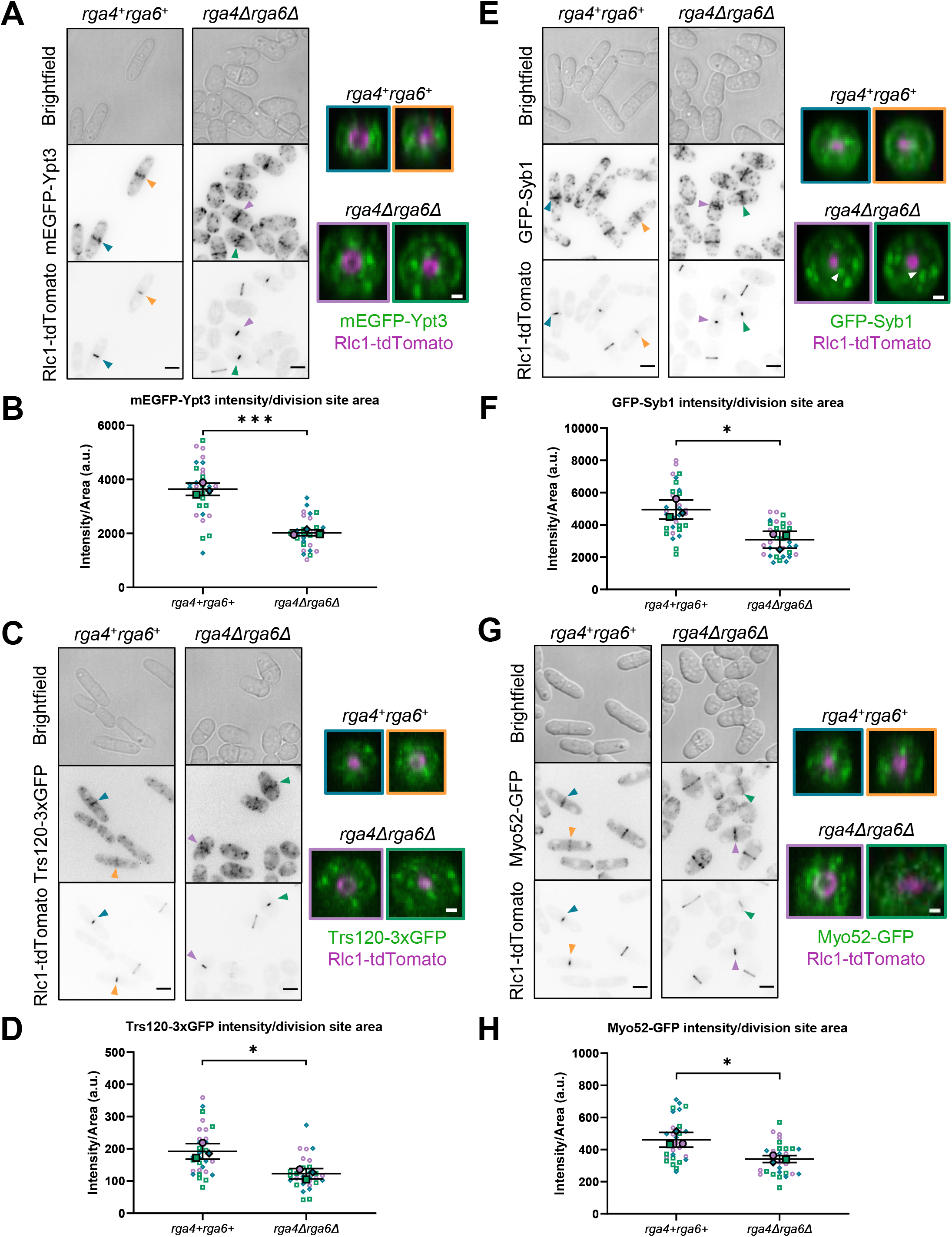
Exocytic markers Ypt3, Syb1, Trs120 and Myo52 do not scale with the enlarged division plane in *rga4Δrga6Δ* mutants. Division site accumulation of exocytic proteins Rab11 GTPase mEGFP-Ypt3 (A-B), TRAPP-II subunit Trs120-3xGFP (C-D), v-SNARE GFP-Syb1 (E-F), and Type V myosin motor Myo52-GFP (G-H) do not scale with increased division site area in the absence of *rga4* and *rga6*; scale bars = 5 μm). Only cells in the late stages of ring constriction were used for this analysis (indicated by arrowheads). 3D division site projections (scale bars = 1 μm) show that each protein besides GFP-Syb1 is properly localized in the absence of *rga4* and *rga6*. GFP-Syb1 is absent from the outer membrane rim in the mutant and instead localizes as bright puncta in the mid-region of the membrane furrow (white arrowheads) (E). 3D-reconstructions are color-coded to arrowheads to indicate cells depicted. Symbol colors in graphs = distinct experiments. Open symbols = individual cells. Solid symbols = means of each experiment. n≥10 cells per genotype per experiment. Unpaired Student’s *t*-test, was used to calculate statistical significance using the means of each experiment. *, *p*< 0.05; ***, *p*<0.001.

We then investigated the localization of the post-Golgi v-SNARE Syb1 which physically tethers exocytic vesicles as a component of the SNARE complex. Similar to Bgs4, Ypt3, and Trs120 during late ring constriction, the sum intensity of GFP-Syb1 normalized to the area of the division site is decreased in the absence of *rga4* and *rga6* (Figure 6F). Notably, the distribution pattern of Syb1 at the division site is slightly altered in *rga4Δrga6Δ* cells. In *rga4*^*+*^*rga6*^*+*^ cells, Syb1 is primarily localized to the outer rim of the division plane. In *rga4Δrga6Δ* mutants, Syb1 at the outer edge of the division plane appears dampened and instead coalesces as large bright puncta throughout the division plane (Figure 6E). Finally, we also examined the localization of the myosin-V motor Myo52, which physically transports exocytic vesicles along actin cables to be delivered throughout the membrane furrow ^6^. After normalizing for division site area, Myo52-GFP sum intensity is less in the absence of *rga4* and *rga6*, similar to the other exocytic proteins examined (Figure 6H). Like the other exocytic proteins aside from Syb1, Myo52 also properly localizes throughout the division plane in the mutant (Figure 6G). In summary, no striking defects in the localization patterns of the exocytic proteins examined were observed, aside from some altered localization of Syb1. Rather than being mislocalized, Ypt3, Trs120, Syb1 and Myo52 appear to be less abundant relative to division site area in the absence of *rga4* and *rga6*. These findings indicate that even though Bgs1 recruitment is enhanced, several vesicular exocytic events do not scale proportionally to the enlarged division site in *rga4Δrga6Δ* mutants.

### Rga4 and Rga6 promotes proper endocytosis

Membrane trafficking includes both delivery of materials via exocytosis as well as recycling of materials via endocytosis. We have previously shown that Cdc42 activity is required for proper endocytic dynamics at the division site ^29^. Here we show that loss of *rga4* and *rga6* results in aberrant membrane organization at the division site. Hence, we asked if excessive Cdc42 activity as seen in *rga4Δrga6Δ* mutants leads to defects in membrane remodeling including endocytosis. To do this, we used the actin filament cross-linker Fim1 to mark sites of endocytosis. Fim1-mEGFP localizes to the division site as observed in sum projections of cells lacking *rga4* and *rga6* (Figure 7A). Next, as with the exocytic markers, we measured the sum intensity of Fim1-mEGFP in both genotypes and then normalized intensity to division site area. After normalization, Fim1-mEGFP sum intensity does not scale to the enlarged division site in the absence of *rga4* and *rga6* (Figure 7B). This indicates that endocytosis is not enhanced in these mutants with respect to the enlarged division plane. Additionally, 3D projections of division sites reveal that unlike in *rga4*^*+*^*rga6*^*+*^ cells, Fim1-mEGFP labeled endocytic patches are not restricted to the outer rim of the membrane furrow but are also visible in the cell interior near the constricting ring in mutant cells (Figure 7C and D).

**Figure 7.**
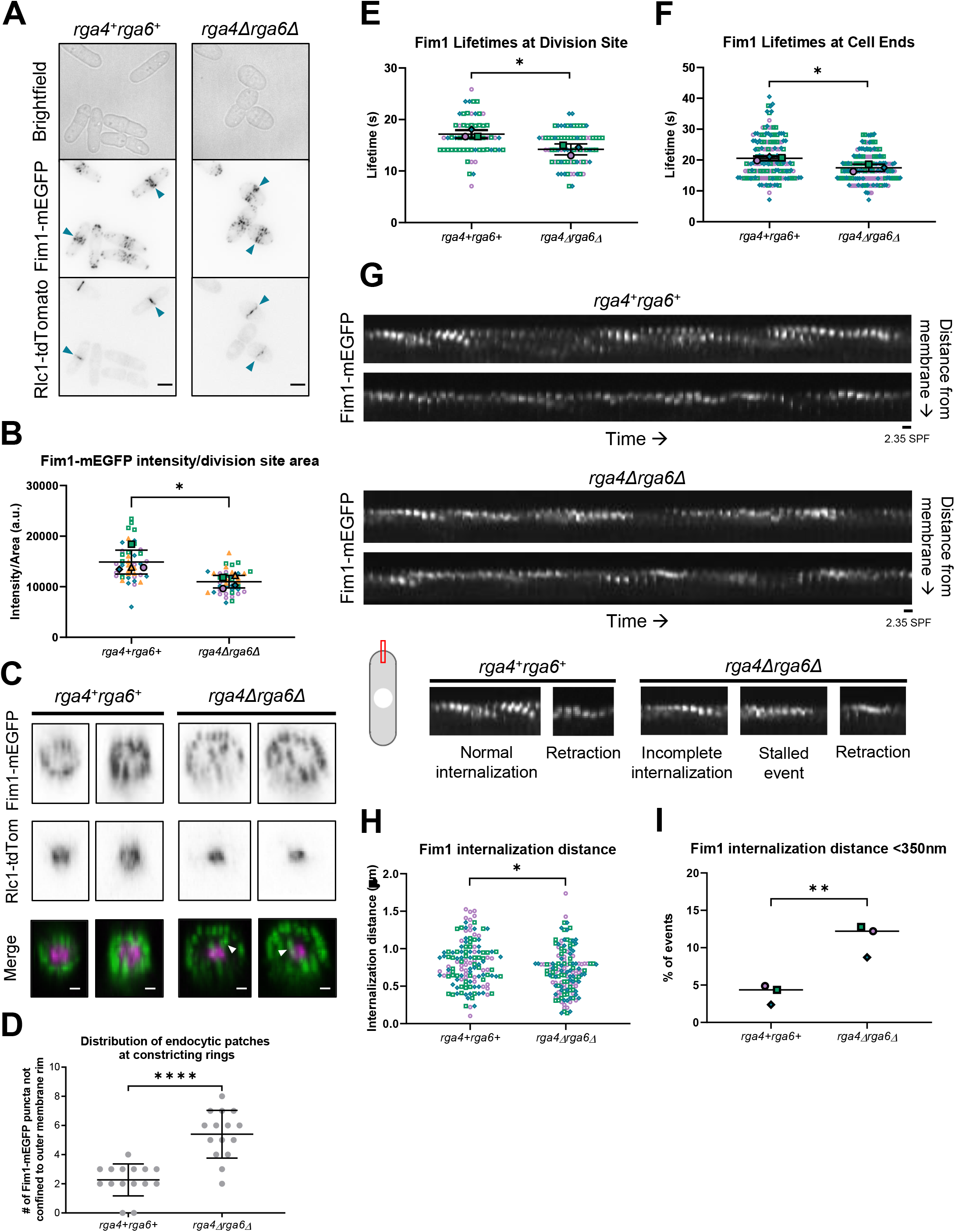
Endocytosis is impaired in the absence of *rga4* and *rga6*. (A) Fim1-mEGP localization at the division site in *rga4*^*+*^*rga6*^*+*^ and *rga4Δrga6Δ* cells (scale bar = 5 μm). (B) Fim1-mEGFP intensity normalized to division site area is less in the absence of *rga4* and *rga6* (n≥10 cells). Only cells in the late stages of ring constriction were used for this analysis (indicated by arrowheads in A). (C) 3D-reconstruction of the division site illustrate that Fim1-marked endocytic patches are not restricted to the outer membrane rim but are also visible near the constricting ring (white arrowheads) in the absence of *rga4* and *rga6* (scale bar = 1 μm). (D) Quantification of C (n=15 cells). (E) At the division site, the lifetimes of Fim1-labeled endocytic patches are shorter on average in the absence of *rga4* and *rga6* (n≥10 endocytic patches per experiment). (F) The lifetimes of Fim1-labeled endocytic patches are also decreased in the absence of *rga4* and *rga6* at the cell ends of interphase cells (n≥45 endocytic patches per experiment). (G) Kymographs generated at the ends of interphase cells illustrate that Fim1-mEGFP labeled endocytic patches do not internalize as far into the cell interior from the plasma membrane in the absence of *rga4* and *rga6*. Additionally, endocytic patches display aberrant motions more frequently in these mutants. These aberrant motions include both stalled and retraction events in which patches fail to properly internalize (scale bar = 800nm; frame rate = 2.35 seconds per frame). (H) Fim1-mEGFP labeled endocytic patches do not internalize as deeply into the cell interior from the plasma membrane in the absence of *rga4* and *rga6* (n≤41 endocytic patches per genotype per experiment). (I) The proportion of endocytic patches that fail to internalize beyond the distance necessary for scission (350 nm) is increased in the absence of *rga4* and *rga6* (n≤41 endocytic patches per genotype per experiment). Symbol colors in graphs = distinct experiments. Open symbols = individual cells. Solid symbols = means of each experiment. Unpaired Student’s *t*-test, was used to calculate statistical significance using the means of each experiment. *, *p*<0.05; **, *p*<0.01; ****, *p*<0.0001.

After observing these defects in the spatial organization of endocytosis in the absence of *rga4* and *rga6*, we then investigated the dynamics of endocytosis in *rga4Δrga6Δ* mutants compared to *rga4*^*+*^*rga6*^*+*^ cells. To do this, we again used Fim1-mEGFP to mark sites of endocytosis and generated time-lapse movies to track the dynamics of the endocytic patches. We find that at the division site the average lifetime of Fim1-mEGFP patches is reduced in *rga4Δrga6Δ* mutants compared to *rga4*^*+*^*rga6*^*+*^ cells (Figure 7E). However, accurately tracking distinct endocytic patches at the division site is challenging due to their dense concentration at that site. Thus, we tracked Fim1-mEGFP labeled endocytic patches at the cell ends of interphase cells instead, using cell polarity to enhance our understanding of cytokinesis as we have used cytokinesis as a paradigm to understand cell polarity ^19,24^. First, we confirmed that the average endocytic patch lifetime is also decreased in the absence of *rga4* and *rga6* at the cell ends (Figure 7F). This suggests that analyzing endocytic patches at the cell ends is a good alternative to the division site. Kymographs of Fim1-mEGFP over time in *rga4*^*+*^*rga6*^*+*^ cells show that the endocytic patches consistently undergo successful internalization (Figure 7G). In contrast, in *rga4Δrga6Δ* mutants we observe an increased incidence of aberrant endocytic patch movements (Figure 7G). These include stalled events in which a patch fails to complete internalization, retraction events in which a patch reverses its motion and moves back to the plasma membrane, and incomplete patch internalization (Figure 7G). We find that overall endocytic patches do not internalize as deeply into the interior of the cell from the plasma membrane in the absence of *rga4* and *rga6* (Figure 7G and H). To clearly illustrate the observed endocytic defect in the absence of *rga4* and *rga6*, we then determined the percentages of endocytic patches which do not internalize beyond 350nm, the distance from the plasma membrane at which vesicle scission is thought to occur ^30^. In *rga4*^*+*^*rga6*^*+*^ cells, approximately 5% of observed endocytic patches fail to internalize beyond 350 nm, while approximately 12% of observed endocytic patches in *rga4Δrga6Δ* cells do not internalize beyond 350 nm suggesting these patches may fail scission (Figure 7I). Together, these observations indicate that loss of *rga4* and *rga6* results in endocytic defects affecting both the distribution of endocytic patches at the division site as well as individual patch dynamics.

## Discussion

Previous reports have shown that membrane trafficking events are essential for successful cytokinesis ^3,6,28^. Our findings illustrate the importance of spatiotemporally regulating membrane trafficking events during cytokinesis to properly coordinate actomyosin ring constriction, membrane furrow ingression, and septum synthesis. In early cytokinesis, Cdc42 promotes the recruitment of Bgs1 that is mainly required for primary septum formation ^2^. Next, Bgs4, that is mainly required for secondary septum formation, is recruited to the division site via a Cdc42-independent mechanism ^2,25^. Delivery of these enzymes via membrane-bound vesicles is coupled to membrane remodeling events necessary for furrow formation ^6^. Here we show that spatiotemporal regulation of Cdc42 activity at the division site helps to maintain proper coupling of the delivery of different septum synthesizing enzymes and membrane remodeling via trafficking events.

The primary septum is built first and only comprises a small portion of the septum, while the secondary septum is formed next and is the major component. This highlights the importance of spatiotemporally regulating the delivery of Bgs1 and Bgs4, the enzymes that synthesize different components of the septum. First, Bgs1 needs to be delivered to the division site where it helps to anchor the actomyosin ring to the membrane and promote primary septum synthesis, thus initiating ring constriction and furrow formation. This is followed by the delivery of Bgs4 at the ingressing membrane to enable secondary septum formation. It is unclear how the delivery of Bgs1 and Bgs4 are regulated to ensure this spatiotemporal distinction. We posit that this is mediated by spatiotemporal regulation of Cdc42 since it is required for the delivery of Bgs1 but not Bgs4. Decreasing Cdc42 activity after onset of septum ingression ensures proper delivery of Bgs4 and corresponding membrane trafficking events. In *rga4Δrga6Δ* mutants, Bgs4 levels do not scale with septum size, which may lead to a delay in septum maturation and a subsequent delay in cell separation. Interestingly, while enhanced Cdc42 activity leads to increased Bgs1 delivery, we did not see any increase in the levels of several exocytic proteins at the division site that are involved in cytokinesis. This indicates that upon misregulation of Cdc42 activity, the balance of the delivery of septum synthesizing enzymes and other membrane trafficking events is disrupted. Further research will mechanistically define these different cargo-specific Cdc42-dependent and -independent delivery events.

Here we also show that misregulation of Cdc42 activity leads to endocytic defects at the division site. Without proper internalization of endocytic vesicles from the division site, the spatial organization of the membrane furrow is likely perturbed. Indeed, we find that *rga4Δrga6Δ* mutants exhibit aberrant septum morphology with membrane trapped within the closed septum. Additionally, the spatial organization of endocytic patches within the membrane furrow is disrupted in the absence of *rga4* and *rga6*. It is possible that this leads to improper internalization of proteins near the constricting ring, thus perturbing the coordination of septum synthesis and membrane furrowing. In the absence of *rga4* and *rga6*, endocytosis is globally misregulated during both cytokinesis and interphase, which suggests that Cdc42 activity must be properly regulated to ensure efficient endocytosis. Indeed, Cdc42 is already known to promote endocytosis when activated by one of its GEF activators, Gef1 ^3^. Interestingly, when Cdc42 activity is impaired in *gef1Δ* cells, an endocytic protein Cdc15 over accumulates in endocytic patches and Cdc15-labeled endocytic patches display longer lifetimes ^3^. Fittingly, these endocytic patch dynamics in *gef1Δ* cells are the exact opposite of those observed in *rga4Δrga6Δ* cells, which indicates that Cdc42 is an important regulator of endocytosis. As a molecular switch, Cdc42 is active when GTP-bound and inactive when bound to GDP. Thus, the molecular regulation of Cdc42 activity may be an efficient way to spatiotemporally control endocytosis to ensure that membrane and proteins are properly recycled from the plasma membrane.

During interphase, active Cdc42 and its GAPs Rga4 and Rga6 display little overlap in their localization since Cdc42 is active at the growing cell ends while these GAPs localize to the cell sides ^21,22^. The other GAP Rga3, conversely, co-localizes with active Cdc42 at growing cell ends as well as at polarity sites in mating cells ^23^. During cytokinesis, however, both active Cdc42 and its GAPs are present at the division site. This observation suggests that some change occurs to minimize the boundary between active Cdc42 and its GAPs during cytokinesis. Recently, Rga4 localization patterns were found to change in a cell cycle-dependent manner ^31^. During interphase, Rga4 localizes to the cell sides in a punctate corset pattern as reported before ^20^, but when cells enter mitosis, Rga4 spreads more homogeneously along the cortex and localizes even up to the cell ends ^31^. Thus, it is possible that cell cycle-dependent regulation of Rga4 enables its localization to the division site to decrease Cdc42 activation after the onset of actomyosin ring constriction. Furthermore, Rga6 displays similar localization dynamics throughout the cell cycle, where it localizes as puncta along the cell sides during interphase, proceeds to the cell ends during early cytokinesis, and then localizes to the division site during ring constriction ^22^. Taken together, here we show that the GAPs Rga4 and Rga6 spatiotemporally regulate Cdc42 activity at the division site to enable proper coupling of membrane furrow ingression and septum synthesis.

## Materials and Methods

### Strains and cell culture

The *S. pombe* strains used in this study are listed in Table 1. All strains are isogenic to the original strain PN567. Cells were cultured in yeast extract (YE) medium and grown exponentially at 25°C. Standard techniques were used for genetic manipulation and analysis ^32^.

### Microscopy

Imaging was performed at room temperature (23–25°C). We used an Olympus IX83 microscope equipped with a VTHawk two-dimensional array laser scanning confocal microscopy system (Visitech International, Sunderland, UK), electron-multiplying charge-coupled device digital camera (Hamamatsu, Hamamatsu City, Japan) and a 100×/1.49 NA UAPO lens (Olympus, Tokyo, Japan). We also used a spinning disk confocal microscope system with a Nikon Eclipse inverted microscope with a 100×/1.49 NA lens, a CSU-22 spinning disk system (Yokogawa Electric Corporation) and a Photometrics EM-CCD camera. Images were acquired with MetaMorph (Molecular Devices, Sunnyvale, CA) and analyzed with ImageJ [National Institutes of Health, Bethesda, MD ^33^]. For still and *z*-series imaging, the cells were mounted directly on glass slides with a #1.5 coverslip (Thermo Fisher Scientific, Waltham, MA) and imaged immediately, and with fresh slides prepared every 10 min. *Z*-series images were acquired with a depth interval of 0.4 μm. For time-lapse images, cells were placed in 3.5 mm glass-bottom culture dishes (MatTek, Ashland, MA) and overlaid with YE medium containing 0.6% agarose and 100 μM ascorbic acid as an antioxidant to minimize toxicity to the cell, as reported previously.

### Electron microscopy

Transmission electron microscopy was performed as described previously (Chappell and Warren, 1989). Cells were washed three times in sterile water, fixed for 1 h in 2% potassium permanganate at room temperature, and then harvested by centrifugation, washed three times in sterile water, resuspended in 70% ethanol, and incubated overnight at 4°C. Samples were then dehydrated by sequential washes in 90% ethanol (twice for 15 min) and washed in 100% ethanol (three times for 20 min). The pellet was resuspended in propylene oxide for 10 min, incubated in a 1:1 mixture of propylene oxide and Spurr’s medium for 1 h, and incubated in neat Spurr’s medium for 1 h. This was followed by another change of medium and incubation at 65°C for 1 h. Finally, samples were embedded in Spurr’s medium in a capsule, and resin in the medium was allowed to polymerize at 60°C overnight. Blocks were sectioned with a diamond knife and stained with uranyl acetate and lead citrate. The cells were then examined in a Zeiss Libra 200MC electron microscope (Oberkochen, Germany) at the University of Tennessee Imaging Core facility.

### Rate of ring constriction analysis

Rate of ring constriction was calculated by dividing the circumference of individual cells by the time each took to complete ring constriction.

### Analysis of fluorescence intensity normalized to division site area

*rga4*^*+*^*rga6*^*+*^ and mutant cells expressing fluorescent proteins were grown to OD 0.2-0.5 and imaged on slides. Depending on the fluorophore, 21-24 *z*-planes were collected at a *z*-interval of 0.4 µm for either or both the 488 nm and 561 nm channels. The same number of z-slices were collected for *rga4*^*+*^*rga6*^*+*^ and mutant cells expressing the same fluorophore using the same imaging settings. ImageJ was used to generate sum projections from the *z*-series and to measure the fluorescence intensity of each fluorophore at the division site. The background fluorescence in a cell-free region of the image or within a dark region inside a cell was subtracted to generate the normalized intensity. The measured intensities were then normalized to the division site area on a cell-by-cell basis. Division site area was calculated using the following formula: A = (3.14 × (D/2)^2^) × 2, in which D = cell diameter (cell width) at the division site. The area was multiplied by 2 in the equation since the division site has two membrane planes, one on each side of the synthesizing septum. The fluorescence intensity of each individual cell measured was then divided by its calculated division site area to determine the amount of fluorescent signal spread across the division site area. A Student’s two-tailed *t*-test, assuming unequal variance, was used to determine significance through comparison of each strain’s mean normalized intensities.

### Analysis of vesicle tracking

The dynamics of endocytic patches marked with Fim1-mEGFP were imaged in a single medial plane with a frame rate of 2.35s for ∼7 minutes. Cells were placed in glass-bottom culture dishes as previously described for time-lapse imaging. To determine endocytic patch lifetime at the division site, Fim1-mEGFP patches were tracked manually. To determine endocytic patch lifetime at the cell ends, Fim1-mEGFP patches were tracked using the Fiji plugin TrackMate where estimated blob diameter = 0.9 micron. Fim1 dynamics were analyzed at the cell ends instead of at the division site since it is easier to spatially resolve distinct patches at the cell ends. Background fluorescence was subtracted from the imaging field away from cells to sharpen the signal of patches for ease of tracking. Lifetime was measured as the time from which a patch first displayed distinguishable fluorescence at the cell cortex to when the fluorescence was no longer detectable. To ensure that patches were tracked from the beginning of patch formation through patch disassembly, patches selected for tracking 1. gradually increased in intensity until a peak intensity was reached and 2. decreased in intensity following peak intensity ^6,34^. Patches tracked with TrackMate were also manually reviewed to ensure tracking accuracy.

Internalization distance of endocytic patches marked with Fim1-mEGFP was measured by generating 5 pixel-wide (800 nm-wide) montages at the cell ends of interphase cells where internalizing patches can be tracked. Again, the cell ends were used for this analysis rather than the division site since it is easier to spatially resolve distinct patches at the cell ends. To measure the distance of internalization, the centroid position of the patch was manually tracked to calculate the displacement of the patch from the cortex as it internalized. The internalization distance was calculated as the distance between the farthest patch centroid position and that of the initial patch centroid position at the cell cortex.

### Statistical tests

GraphPad Prism was used to determine significance. One-way ANOVA, followed by a Tukey’s multiple comparisons post-hoc test, was used to determine individual *p*-values when comparing three or more samples. When comparing two samples, an unpaired Student’s *t*-test (two-tailed, unequal variance) was used to determine significance.

## Acknowledgments

We thank Dr. Andreas Nebenführ for discussions; Dr. Fred Chang, Dr. Pilar Perez, Dr. Sophie Martin, Dr. Fulvia Verde, and Dr. Jian-Qiu Wu for strains; and Dr. John Dunlap for electron microscopy. This research was funded by the National Science Foundation CAREER award #1941367 to M.D. The authors do not have any conflict of interest.

