## Supplemental Figures for "Cdc42 GTPase activating proteins Rga4 and Rga6 coordinate septum synthesis and membrane trafficking at the division plane during cytokinesis"

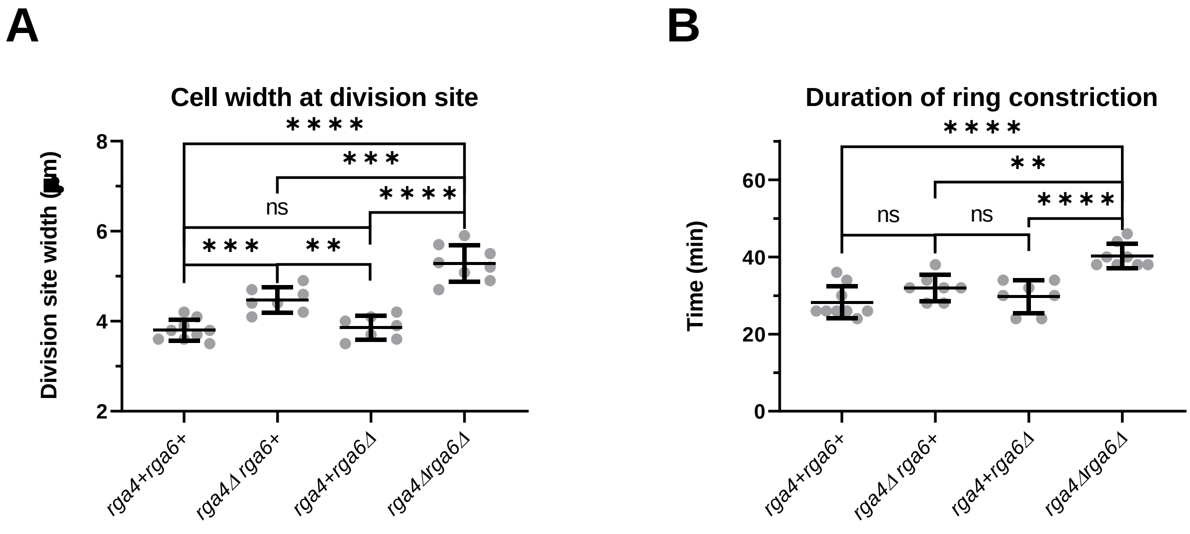


**Supplemental Figure S1.** Loss of *rga4* and *rga6* results in wider cells. (A) Loss of *rga4* and *rga6* additively increases cell width compared to *rga4*$^{+}$*rga6*$^{+}$ cells. (B) As wider cells with larger actomyosin rings, *rga4Δrga6Δ* cells take longer to constrict their rings relative to *rga4*$^{+}$*rga6*$^{+}$ and the single GAP mutants. Statistical test used is Ordinary one-way ANOVA with Tukey's multiple comparisons. n.s., not significant; *, $p< 0.05$; **, $p< 0.01$; ***, $p<0.001$; ****, $p< 0.0001$.

### Supplemental Figure S1

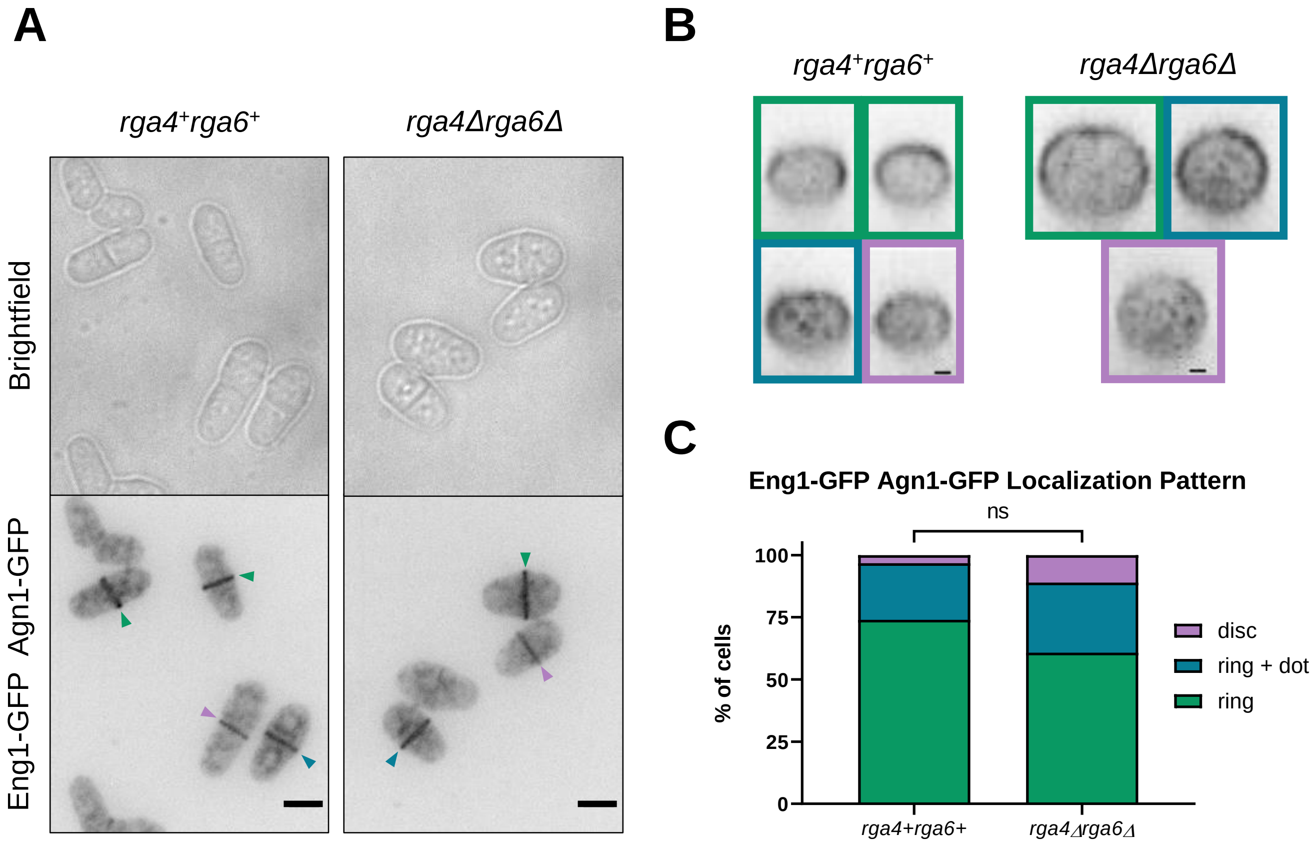


**Supplemental Figure S2.** The primary septum digesting glucanases Eng1 and Agn1 localize properly in the absence of *rga4* and *rga6*. (A) Eng1-GFP and Agn1-GFP localize to the division site in the absence of *rga4* and *rga6* in a manner similar to *rga4*$^{+}$*rga6*$^{+}$ cells (blue, green, and purple arrows; scale bar = 5$\mu$m). (B) 3D-reconstructions of the division sites of *rga4*$^{+}$*rga6*$^{+}$ and *rga4Δrga6Δ* cells illustrate that Eng1-GFP and Agn1-GFP properly localize to both the outer rim of the membrane furrow as a ring (green panels corresponding to cells marked with green arrowheads) and to the center of the division plane as a dot in addition to the membrane rim (blue panels corresponding to cells marked with blue arrowheads). Few cells in either genotype exhibited mislocalization of Eng1-GFP and Agn1-GFP as a disk (purple panels corresponding to cells marked with purple arrowheads; scale bar = 1$\mu$m). (C) Quantification of Eng1-GFP and Agn1-GFP localization patterns at the division site in *rga4*$^{+}$*rga6*$^{+}$ and *rga4Δrga6Δ* cells, where N = 3 replicate experiments, $n \geq 29$ cells for each genotype. Statistical test used was unpaired Student's *t*-test, n.s., not significant.

### Supplemental Figure S2

**Table 1.** List of strains used in this study

| Strain | Genotype | Source |
|---|---|---|
| PN567 | h+ ade6-704 leu1-32 ura4-d18 | Paul Nurse |
| FV513 | h- Δrga4::ura4+ ade6-704 leu1-32 ura4-D18 | Das et al., 2007 |
| PPG45.11 | h+ Δrga6::kanMX ade- leu1-32 ura4-D18 | Revilla-Guarinos et al., 2016 |
| YMD672 | h- Δrga4::ura4+ Δrga6::kanMX ade- leu1-32 ura4-D18 | This study |
| YMD314 | CRIB-3xGFP-ura4+ rlc1-Tomato-NATr sad1-mCherry:kanMX ade6-M21X leu1-32 ura4-D18 | Lab stock |
| YMD746 | Δrga6::kanMX CRIB-3xGFP-ura4+ rlc1-Tomato-NATr sad1-mCherry:kanMX ade6-M21X leu1-32 ura4-D18 | This study |
| YMD754 | Δrga4::kanMX CRIB-3xGFP-ura4+ rlc1-Tomato-NATr sad1-mCherry:kanMX ade6-M21X leu1-32 ura4-D18 | This study |
| YMD787 | Δrga4::ura4+ Δrga6::kanMX CRIB-3xGFP-ura4+ rlc1-Tomato-NATr sad1-mCherry:kanMX ade6-M21X leu1-32 ura4-D18 his7+ | This study |
| YMD771 | rga6-GFP-kanMX rlc1-tdTomato-NATr sad1- mCherry:kanMX ade6-M216 leu1-32 ura4-D18 | This study |
| YMD772 | rga4-GFP-kanMX rlc1-tdTomato-NATr sad1- mCherry:kanR ade6-M216 leu1-32 ura4-D18 | This study |
| YMD1865 | rga3-GFP-kanMX rlc1-tdTomato-NATr ade6-M216 leu1-32 ura4-D18 | This study |
| YMD1463 | eng1-GFP-kanMX agn1-GFP-kanMX rlc1-tdTomato-NATr ade6-M216 leu1-32 ura4-D18 | This study |
| YMD1465 | Δrga4::ura4+ Δrga6::kanMX eng1-GFP-KanMX agn1-GFP-KanMX rlc1-tdTomato-NATr ade6-M216leu1-32 ura4-D18 | This study |
| YMD1452 | Δbgs1::ura4 Pbgs1::GFP-bgs1:leu1+ rlc1-tdTomato-NATr$^r$ leu1-32 ura4-D18 | Onwubiko et al., 2020 |
| YMD1848 | Δrga4::ura4+ Δrga6::kanMX Δbgs1::ura4 Pbgs1::GFP-bgs1:leu1+ rlc1-tdTomato-NATr$^r$ leu1-32 ura4-D18 | This study |
| YMD1447 | h- Δbgs4::ura4 Pbgs4::GFP-bgs4:leu1+ rlc1-tdTomato-NATr$^r$ sad1-mCherry:kanMX leu1-32 ura4-D18 | This study |
| YMD1449 | Δrga4::ura4+ Δrga6::kanMX Δbgs4::ura4 Pbgs4::GFP-bgs4:leu1+ rlc1-tdTomato-NATr$^r$ sad1-mCherry:kanMX leu1-32 ura4-D18 | This study |
| YMD1670 | kanMX6-Pypt3-mEGFP-ypt3 rlc1-tdTomato-NATr ade6-210 leu1-32 ura4-D18 | This study |
| YMD1711 | Δrga4::ura4+ Δrga6::kanMX kanMX6-Pypt3-mEGFP-ypt3 rlc1-tdTomato-NATr ade6-210 leu1-32 ura4-D18 | This study |
| JW6731-1 | h- trs120-3GFP-kanMX6 rlc1-tdTomato-natMX6 ade6-M210 leu1-32 ura4-D18 | Wang et al., 2016 |
| YMD1679 | Δrga4::ura4+ Δrga6::kanMX trs120-3GFP-kanMX6 rlc1-tdTomato-natMX6 ade6-M210 leu1-32 ura4-D18 | This study |
| YMD1751 | kanMX-GFP-syb1 rlc1-tdTomato-natMX6 ade6-M216 leu1-32 ura4-D18 | This study |
| YMD1755 | Δrga4::ura4+ Δrga6::kanMX kanMX-GFP-syb1 rlc1-tdTomato-natMX6 ade6-M216 leu1-32 ura4-D18 | This study |
| YMD1756 | myo52-GFP::kanMX rlc1-tdTomato-natMX6 ade6-M216 leu1-32 ura4-D18 | This study |
| YMD1731 | Δrga4::ura4+ Δrga6::kanMX myo52-GFP::kanMX rlc1-tdTomato-natMX6 ade6-M216 leu1-32 ura4-D18 | This study |
| JW6771-1 | h+ fim1-mEGFP-kanMX6 rlc1-tdTomato-natMX6 ade6-M210 leu1-32 ura4-D18 | Wang et al., 2016 |
| YMD1110 | Δrga4::ura4+ Δrga6::kanMX fim1-mEGFP-kanMX6 rlc1-tdTomato-natMX6 ade6-M210 leu1-32 ura4-D18 | This study |
